# On a Unifying ‘Reverse’ Regression for Robust Association Studies and Allele Frequency Estimation with Related Individuals

**DOI:** 10.1101/470328

**Authors:** Lin Zhang, Lei Sun

## Abstract

For genetic association studies with related individuals, standard linear mixed-effect model is the most popular approach. The model treats a complex trait (phenotype) as the response variable while a genetic variant (genotype) as a covariate. An alternative approach is to reverse the roles of phenotype and genotype. This class of tests includes quasi-likelihood based score tests. In this work, after reviewing these existing methods, we propose a general, unifying ‘reverse’ regression framework. We then show that the proposed method can also explicitly adjust for potential departure from Hardy–Weinberg equilibrium. Lastly, we demonstrate the additional flexibility of the proposed model on allele frequency estimation, as well as its connection with earlier work of best linear unbiased allele-frequency estimator. We conclude the paper with supporting evidence from simulation and application studies.

## 1 Introduction

Genetic association studies aim at identifying genetic variants, *G*s, that influence a heritable trait, *Y*, of interest. To this end, allele-based association tests or allelic tests, comparing allele frequencies between case and control groups, are locally most powerful (Sasieni, 1997) in a sample of unrelated individuals. However, traditional allelic tests (i) analyze only binary outcomes, (ii) cannot easily accommodate covariates, (iii) are limited to independent samples, and (iv) have type I error control issue if there is a departure from Hardy–Weinberg equilibrium (HWE) in the study population.

HWE states that the two alleles in a genotype are independent draws from the same distribution, or, equivalently, genotype frequencies depend solely on the allele frequencies. For a bi-allelic SNP with two possible alleles *A* and *a*, let *p* and 1 − *p* be the respective allele frequencies. Under HWE, *p*_*aa*_ = (1 − *p*)^2^, *p*_*Aa*_ = 2*p*(1 − *p*), and *p*_*AA*_ = *p*^2^, where *p*_*aa*_, *p*_*Aa*_, and *p*_*AA*_ are the genotype frequencies of genotypes *aa, Aa* and *AA*, respectively. To measure the departure from HWE or the amount of Hardy–Weinberg disequilibrium (HWD),

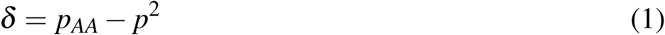

is a widely used quantity, and *δ* = 0 corresponds to HWE (Weir, 1996).

Genotype-based association tests treat phenotype *Y* as the response variable and genotype *G* as an explanatory variable. Due to the regression nature of the framework, genotype-based association tests can easily handle continuous traits and incorporate covariates. It is commonly assumed that genotype-based association tests are robust to departure from HWE. With a sample of independent individuals, both theoretical and empirical results support this (Sasieni, 1997; Schaid and Jacobsen, 1999). However, in the presence of sample dependency, little has been discussed.

When individuals in a sample are genetically related with each other, linear mixed-effect models (LMM) have become the most popular approach for association testing (Eu-Ahsunthornwattana et al., 2014). The variance-covariance matrix of the phenotype is partitioned into a weighted sum of correlation structure due to genetic relatedness and shared environmental effects, where the weight is usually referred to as ‘heritability’ (Visscher et al., 2008). The genetic relatedness is typically represented by a known kinship coefficient matrix, or estimated based on the available genome-wide genetic data if the pedigree information was not collected (Yang et al., 2011; Sun and Dimitromanolakis, 2012).

An alternative approach is to reverse the roles of *Y* and *G* in the regression model. O’Reilly et al. (2012) proposed MultiPhen, a method that treats the genotype *G* of a SNP as the response variable and phenotype values *Y* s of multiple traits as predictors. However, MultiPhen relies on ordinal logistic regression and analyzes only independent samples. MultiPhen does not require the assumption of HWE, but insights to its robustness to HWD was not provided (O’Reilly et al., 2012).

Thornton and McPeek (2007) extended the traditional allelic tests to study binary traits with related individuals. Their test was then generalized by Feng et al. (2011) and Feng (2014) to a quasi-likelihood score test for either binary or continuous traits. However, none of these methods can directly incorporate covariates. Jakobsdottir and McPeek (2013) later proposed a ‘retrospective’ approach, MASTOR, to study the association between *G* and one (approximately) normally distributed trait *Y*, while accommodating covariates in related individuals. All methods in this category, however, implicitly assumed HWE.

In this paper, we first review and provide some insights into the aforementioned genetic association tests. We then propose a robust and flexible ‘reverse’ regression framework that (a) unifies several existing association methods, and (b) explicitly includes a correction factor in the variance-covariance matrix to adjust for potential departure from HWE. Further, we show that the proposed ‘reverse’ regression framework (c) can also be used to estimate allele frequency in complex pedigrees. Interestingly, we reveal that for the simple case of no covariates and HWE, the proposed estimator is the best linear unbiased estimator of McPeek et al. (2004). We conclude the paper with supporting evidence from simulation and application studies, and some discussion points.

## 2 Method

### 2.1 The Traditional Allele-based Association Test, or Allelic Test, *T*_allelic_

Consider a bi-allelic SNP with two alleles *a* and *A*. Without loss of generality, *A* is the minor allele with population minor allele frequency (MAF) less than 0.5. Consider a case-control study with independent observations. We use *r*_*k*_, for *k* ∈ {0, 1, 2}, to denote the genotype counts, respectively, for genotypes *aa, Aa* and *AA*, in the case group of size *r*. Similarly, *s*_*k*_ for the control group of size *s*, and *n*_*k*_ for the combined sample of size *n*.

Let 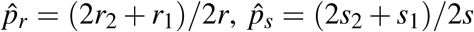, and 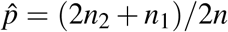 be the sample allele frequencies of allele *A*, respectively, in the case, control and combined samples. The classical allelic association test is based on,

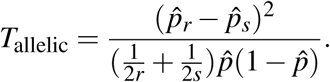

Under the null of no association, 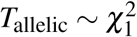.

It has been shown that *T*_allelic_ is locally most powerful for a sample of unrelated individuals (Sasieni, 1997). However, the validity of *T*_allelic_ requires the assumption of Hardy–Weinberg equilibrium. Some remedies have been proposed. For example, Schaid and Jacobsen (1999) considered a variance adjustment and recommended 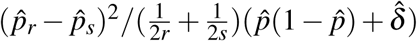 as a robust allelic test, where

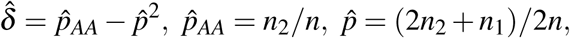

and 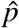 and 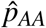 are the sample frequency estimates of allele *A* and genotype *AA*, respectively. Nevertheless, existing (classical and robust) allelic tests are limited to binary *Y* without consideration of covariate effects.

### 2.2 The Traditional Phenotype-on-Genotype (*Y* -on-*G*) Association Tests, *T*_indep_ and *T*_LMM_

#### 2.2.1 Independent Samples, *T*_indep_

Define *G* = 0, 1 and 2 for genotypes *aa, Aa* and *AA*, respectively, and let *Y* be a (continuous) trait of interest. With a sample of *n* unrelated individuals, the traditional genotype-based association test assumes that,

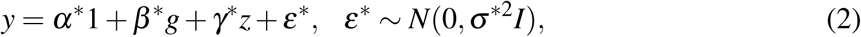

where *y* = (*y*_1_, *y*_2_, …, *y*_*n*_) is a *n* × 1 vector for the phenotypic values, *g* = (*g*_1_, *g*_2_, …, *g*_*n*_) is a *n* × 1 vector for the genotypes of the SNP, *ε*\* is the error term with variance *σ*\*^2^, 1 is a *n* × 1 vector of 1s, and *I* is the identity matrix. For notation simplicity but without loss of generality, we assume that there is only one additional covariate, denoted by *z*.

Score tests are often used for genetic association analyses (Derkach et al., 2015). In this case, the score statistic of testing *H*_0_ : *β*\* = 0 can be easily derived as

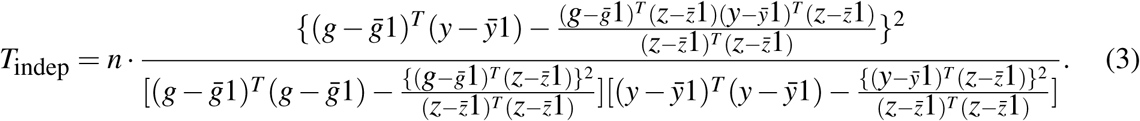

After some simple algebraic manipulations, one can show that

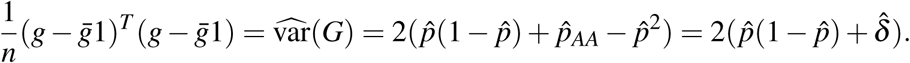

Because 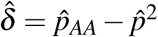 measures the amount of Hardy–Weinberg disequilibrium present in the data (Weir (1996)), *T*_indep_ inherently adjusts for departure from HWE through 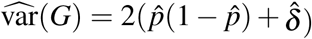. As a result, the traditional genotype-based association test is robust to HWD in independent samples.

When *Y* is binary, logistic regression is commonly used. However, Chen (1983) showed that under common regularity conditions, the score test statistic takes identical form for exponential family in independent samples with no covariates. (Derkach et al. (2015) also showed that for *Y* -dependent sampling, “the score statistics are identical for conditional and full likelihood approaches, and are of the same form as for ordinary random sampling.”) Thus, in terms of association testing, we can conclude that genotype-based association studies of binary traits in independent samples are also robust to departure from HWE.

#### 2.2.2 Dependent Samples, *T*_LMM_

When individuals in a sample are related to each other as in pedigree data, a common practice is to replace var(*ε*\*) = *σ*\*^2^*I* in (2) with 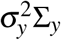 to reflect the sample dependence, and use the linear mixed-effect model,

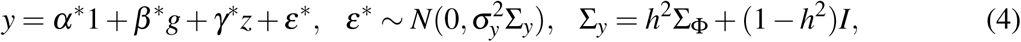

where *h*^2^ is the ‘heritability’, and Σ_Φ_ is the kinship coefficient matrix. Among the total variance of 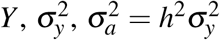 can be interpreted as the variance of *Y* due to additive genetic variation, while 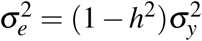 as the variance due to environmental variation.

For the Σ_Φ_ matrix, Σ_Φ_(*i, j*) = 2*Φ*_*i,j*_ where *Φ*_*i,j*_ is the kinship coefficient between individual *i* and individual *j*. When the pedigree information is not available, it is a common practice to estimate Σ_Φ_(*i, j*) with the averaged sample correlation across *K* autosomal SNPs (Yang et al., 2011),

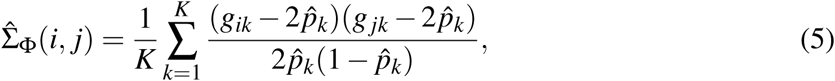

where *g*_*ik*_ and *g* _*jk*_ are the genotypes of SNP *k* for individual *i* and *j* respectively, and 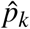 is the estimated allele frequency.

Without the assumption of HWE, var(*G*_*k*_) = 2(*p*_*k*_(1 − *p*_*k*_) + *δ*_*k*_). Thus, (5) implies *δ*_*k*_ = 0, and the traditional kinship coefficient estimation implicitly assumes that HWE holds at each and every SNP *k, k* = 1, …, *K*. Consequently, *T*_LMM_ derived from the linear mixed model (4) can be sensitive to departure from HWE. In Section 3, we will demonstrate with a simple sib-pair design that whenthe true heritability is known, the empirical type I error rate of the standard linear mixed-effect model is inflated when *δ* > 0 (or deflated if *δ <* 0), and when heritability is estimated from the data, the test is accurate but the estimated heritability is then biased when *δ* ≠ 0.

### 2.3 The Proposed ‘Reverse’ Genotype-on-Phenotype (*G*-on-*Y*) Regression Model and Association Test, *T*_reverse_

To account for potential departure from Hardy–Weinberg equilibrium yet adjusting for covariate effect and sample dependency, one useful direction is to reverse the roles of *Y* and *G* in the regression model. We define the ‘reverse’ regression model as,

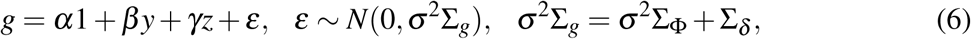

where Σ_Φ_ is the kinship coefficient matrix as defined earlier, and Σ_*δ*_ is a function of *δ*, that explicitly models the amount of Hardy–Weinberg disequilibrium. For example, for a pair of siblings,

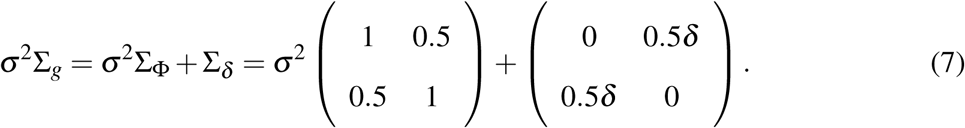

Under the null hypothesis of *H*_0_ : *β* = 0, the corresponding score test statistic is

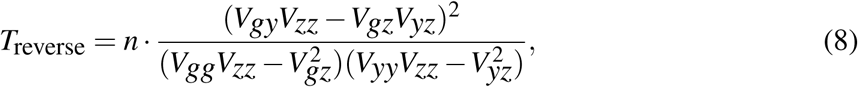

where

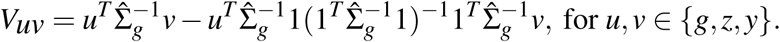

*T*_reverse_ is asymptotically 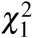 distributed under *H*_0_.

The response variable *G*, in the proposed ‘reverse’ regression (6), is a discrete random variable with three possible values 0, 1, and 2. However, in light of the result of Chen (1983), we modelled *G* using a linear model. We show in the following that indeed, in the simple case of independent sample, HWE, or no covariates, *T*_reverse_ derived from (6) shares similar properties with previous methods that treat *G* as discrete.

### 2.4 Connections with *T*_indep_ and *T*_LMM_

In the absence of sample correlation, Σ_*g*_ = *I*, and the proposed test statistic in (8) is then simplified to *T*_indep_ in (3); *T*_reverse_ is symmetric with respect to *G* and *Y*, provided that the same Σ is used. Thus, for independent samples, *T*_reverse_ adjusts for HWD through var(*G*) = 2(*p*(1 − *p*) + *δ*) as in the traditional association test, and

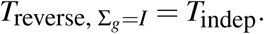

In the presence of sample correlation. it is not difficult to see that if both *T* _LMM_ and *T*_reverse_ were to use the same, known correlation matrix Σ in their respective regression models (4) and (6), then the two tests are equivalent with each other,

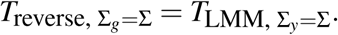

However, the traditional linear mixed-effect model uses kinship coefficient matrix Σ_Φ_ to model Σ_*y*_, while the proposed method adjusts for sample correlation through Σ_*g*_ that includes both Σ_Φ_ for relatedness and a correction factor *δ* in Σ_*δ*_ for potential HWD at the tested SNP.

### 2.5 Connections with Quasi-likelihood Association Tests without Covariates, *T*_QL-binary_ and *T*_GQLS_

For case-control studies with related samples, Thornton and McPeek (2007) proposed an association test that compares sample allele frequency estimates between the case and control groups, while adjusting for relatedness between individuals. The test statistic is defined as,

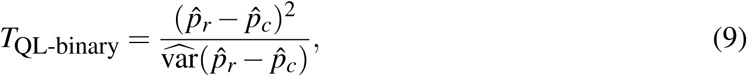

where

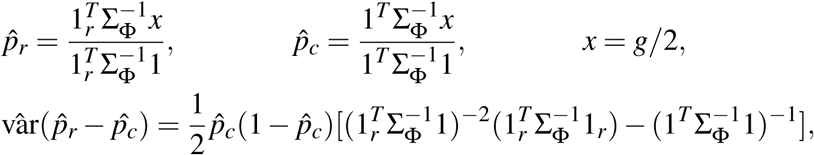

and 1_*r*_ is a *n* × 1 vector with the *i*th observation to be 1 if individual *i* is from the case group, and 0 otherwise. Note that 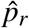 is a sample estimate of the allele frequency in cases, while 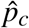 is the pooled estimate using the case-control combined sample, both adjusting for sample correlation through Σ_Φ_.

Feng et al. (2011) and Feng (2014) reformulated *T*_QL-binary_ as a generalized quasi-likelihood score test that handles both continuous and binary traits but does not allow for covariates. The quasi-likelihood based regression framework assumes that,

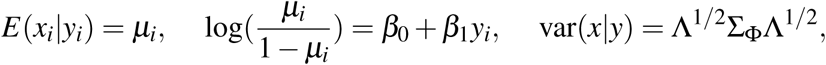

where Λ is an *n*×*n* diagonal matrix, and {diag{Λ}} = {*µ*_1_(1 − *µ*_1_), *µ*_2_(1 − *µ*_2_), …, *µ*_*n*_(1 − *µ*_*n*_)}. Under the null of no association, the framework also assumes that *E*(*x*_*i*_) = *p*_*c*_, and var(*x*) = *p*_*c*_(1 − *p*_*c*_)Σ_Φ_. Thus, the generalized quasi-score statistic of testing *H*_0_ : *β*_1_ = 0 is,

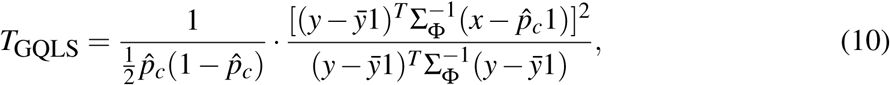

where 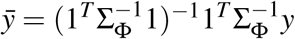, and 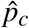 is identical to that in *T*_QL-binary_ of (9). When *Y* is binary, *y* = 1_*r*_. Substituting *y* with 1_*r*_ in (10), it is easy to show that

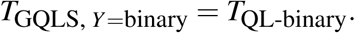

When individuals are independent of each other in a sample, Σ_Φ_ = *I*, then 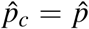. Thus, 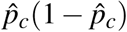 in the denominator of (10) assumes *δ* = 0 (HWE) in estimating var(*G/*2). In contrast,

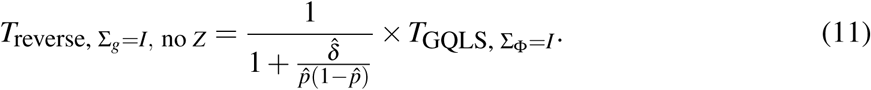

### 2.6 Connections with MASTOR, an Alternative Mixed-Model Association Test with Covariates, *T*_MASTOR_

For continuous traits with covariates, Jakobsdottir and McPeek (2013) proposed MASTOR, a mixed-effect model-based association test. The MASTOR test statistic of no association is defined as,

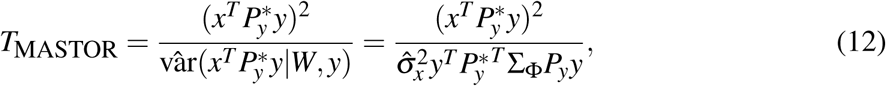

where

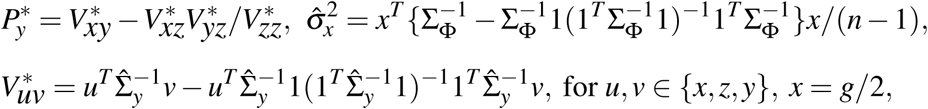

and Σ_*y*_ is the same as that in the linear mixed-effect model of (4), and 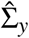 is the maximum likelihood estimate under the null of no association.

To compare *T*_reverse_ and *T*_MASTOR_, we rewrite *T*_reverse_ in (8) as

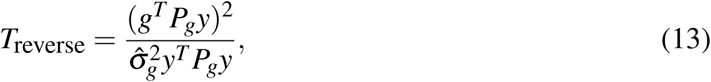

where 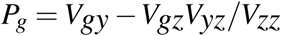, and 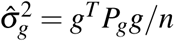.

Note that *V*_*uv*_ in (8) (or re-formulated as in (13)) has the same form as 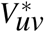 above. Thus, if we were to use Σ_*y*_ = Σ_Φ_ in the calculation of 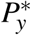 for *T*_MASTOR_ (i.e. assume *h*^2^ = 1 in Σ_*y*_ = *h*^2^Σ_Φ_ + (1 *-h*^2^)*I*), and not accounting for HWD in the calculation of *T*_reverse_ (i.e. assume *δ* = 0 in *Σ* ^2^Σ_Φ_ + Σ_*δ*_), then

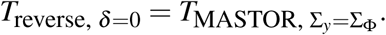

However, although both *T*_MASTOR_ and *T*_reverse_ measure the correlation between *G* and *Y* while adjusting for covariate effects, the two approaches are distinct from each other. *T*_reverse_ in (8) is directly derived from a regression framework with *G* as the response variable and *Y* as a covariate, explicitly adjusting for HWD through *Σ* ^2^Σ_*g*_ = *Σ* ^2^Σ_Φ_ + Σ_*δ*_ for the tested SNP using model (6). That is, *V*_*uv*_ uses Σ_*g*_ that contains both Σ_Φ_ and Σ_*δ*_ as illustrated in (7) for sibling pairs. In contrast, *T*_MASTOR_ in (12) is a hybrid score statistic, where the score function is derived from the linear mixed-effect model with *Y* as the response variable, while the variance of the score function is estimated from a ‘retrospective’ approach that models *X* = *G/*2 as the response variable. When estimating 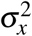 and modelling Σ_*y*_, *T*_MASTOR_ considers Σ_Φ_ alone to account for the genetic relationship between related individuals. Hence, *T*_MASTOR_ implicitly assumes HWE.

### 2.7 Allele Frequency Estimation with the Proposed ‘Reverse’ Regression

Another feature of the proposed ‘reverse’ regression framework is its inherent ability to estimate allele frequencies in dependent samples while adjusting for covariate effects. When no phenotype *Y* is included in (6), the ‘reverse’ regression is simplified to

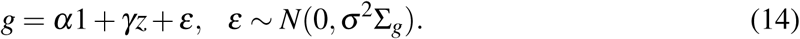

The intercept *α* can be interpreted as twice the population allele frequency, *p*, for the SNP under the study. The maximum likelihood estimator of *α* is

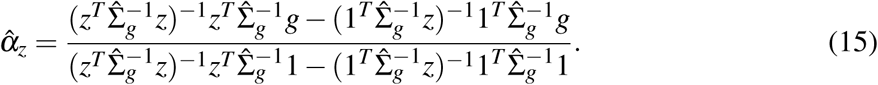

When no covariate is included in (14) and Σ_*g*_ is replaced with the kinship coefficient matrix 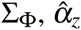 is then reduced to

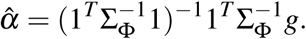

Interestingly, 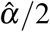 is the best linear unbiased estimator of *p* as studied in McPeek et al. (2004). Thus, the proposed 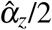 is a generalized allele frequency estimator that, in addition to accounting for sample dependency, can also adjust for covariate effects and potential departure from Hardy– Weinberg equilibrium.

## 3 Empirical Results

Given the analytical insights provided above, here we briefly examine the performance of the commonly used *T*_LMM_ based on the linear mixed-effect model of (4) and the proposed *T*_reverse_ based on (6) for association analyses, through application and simulation studies. We also demonstrate numerically the additional utility of *T*_reverse_ for allele frequency estimation.

### 3.1 Cystic Fibrosis Sib-pair Data

We first extracted 65 sibling pairs from a cystic fibrosis (CF) gene modifier study, as previously described in Sun et al. (2012) and Wright et al. (2011). The outcome *Y* of interest is the lung function measurements of the 130 CF subjects. In total, there were 570, 539 SNPs genotyped using the Illumina 610-Quad Beadchip. To stabilize the variance estimation, we also required SNPs to have minor allele frequency greater than 5%. We then applied the *T*_LMM_ and the proposed *T*_reverse_ to the remaining 505, 172 SNPs. For the implementation of *T*_LMM_, we treated *h*^2^ as unknown and estimated it based on the linear mixed-effect model of (4) as in convention.

As expected, the results of *T*_LMM_ and the proposed *T*_reverse_ are similar to each other (Figure 1), and both have good type I error control (results not shown). However, the estimated *h*^2^, obtained using the 65 sibling data, is 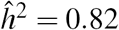. This value is substantially greater than 0.5, the commonly-believed ‘true’ heritability of lung function in CF (Vanscoy et al., 2007). To verify if the biased heritability estimate is due to chance, we conducted a simulation study assuming that only one causal SNP, *G*_causal_ with MAF of 0.2, affects *Y* with *h*^2^ = 0.5. Genotype and phenotype values for 65 sibling pairs were then simulated under the assumption of HWE. Among the 100, 000 independently generated replicates, only 4.24% of the heritability estimates was greater than 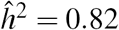, the value that was observed in the real CF data.

**Figure 1:**
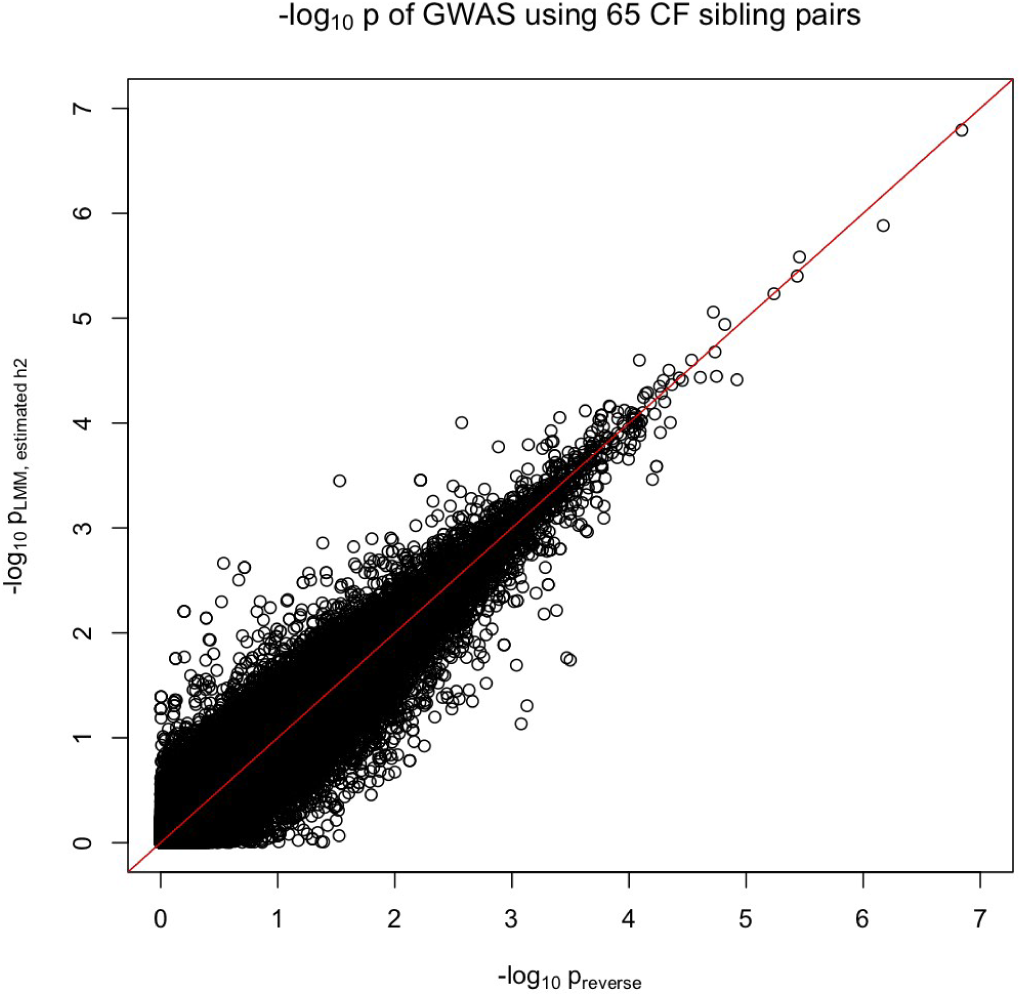
CF sib-pair data application results, comparing *T*^LMM^ based on the linear mixed-effect model (4) and *T*_reverse_ based on the proposed ‘reverse’ regression model (6). The association is between lung function measurements in 65 sibling paris with cystic fibrosis and 505, 172 autosomal SNPs (with minor allele frequency greater than 0.05). Implementation of *T*_LMM_ assumed *h*^2^ unknown, and 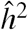 estimated based on 65 samples was 0.82; heritability of lung function in CF is believed to be around 0.5 based on CF epidemiological studies (Vanscoy et al., 2007).

One possible explanation is that the causal SNP(s) may be not be in HWE, resulting in a biased estimator of heritability, 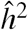 . Following the same sib-pair design, we next used simulations to demonstrate that (a) assuming the true *h*^2^ is known, the empirical type I error rate of linear mixed-effect model (4) inflates if *δ* > 0, or deflates if *δ* < 0, and (b) when *h*^2^ is treated as a nuisance parameter, its estimate based on model (4) can be biased in the presence of Hardy– Weinberg disequilibrium.

### 3.2 Simulated Sib-pair Data

Consider a continuous trait *Y* with *h*^2^ = 0.5 and influenced by one causal SNP with minor allele frequency of 0.2, *G*_causal_, for which the HWD factor is *δ*_causal_. We conducted association testing between *Y* and a non-associated SNP, *G*_tested_ (with its own *δ*_tested_) using simulated data from 65 sibling pairs. The sample size 65 was chosen to match with the number of sibling pairs in the cystic fibrosis dataset in Section 3.1.

Figure 2(a) plots the empirical type I error rates (black circles) of *T*_LMM_ using the true *h*^2^ = 0.5, for a nominal level of 0.05, estimated from independently simulated 10, 000 replicates for each *δ*_causal_ value. The trend of type I error inflation is clear as *δ*_causal_ increases. In Figure 2(a) we set *δ*_tested_ = 0.06, but we note that the root cause of the type I error issue is *δ*_causal_ ≠ 0 when using the LMM (4). Figure 2(b) shows that even if *G*_tested_ is in HWE (*δ*_tested_ = 0) the problem remains, albeit less severe, as long as *δ*_causal_ ≠ 0. The simulation results also confirm that the proposed *T*_reverse_ has the correct type I error control (red triangles in Figure 2); results are similar if *G*_tested_ were not simulated but based on the real genotype data observed in the 65 sibling pairs.

**Figure 2:**
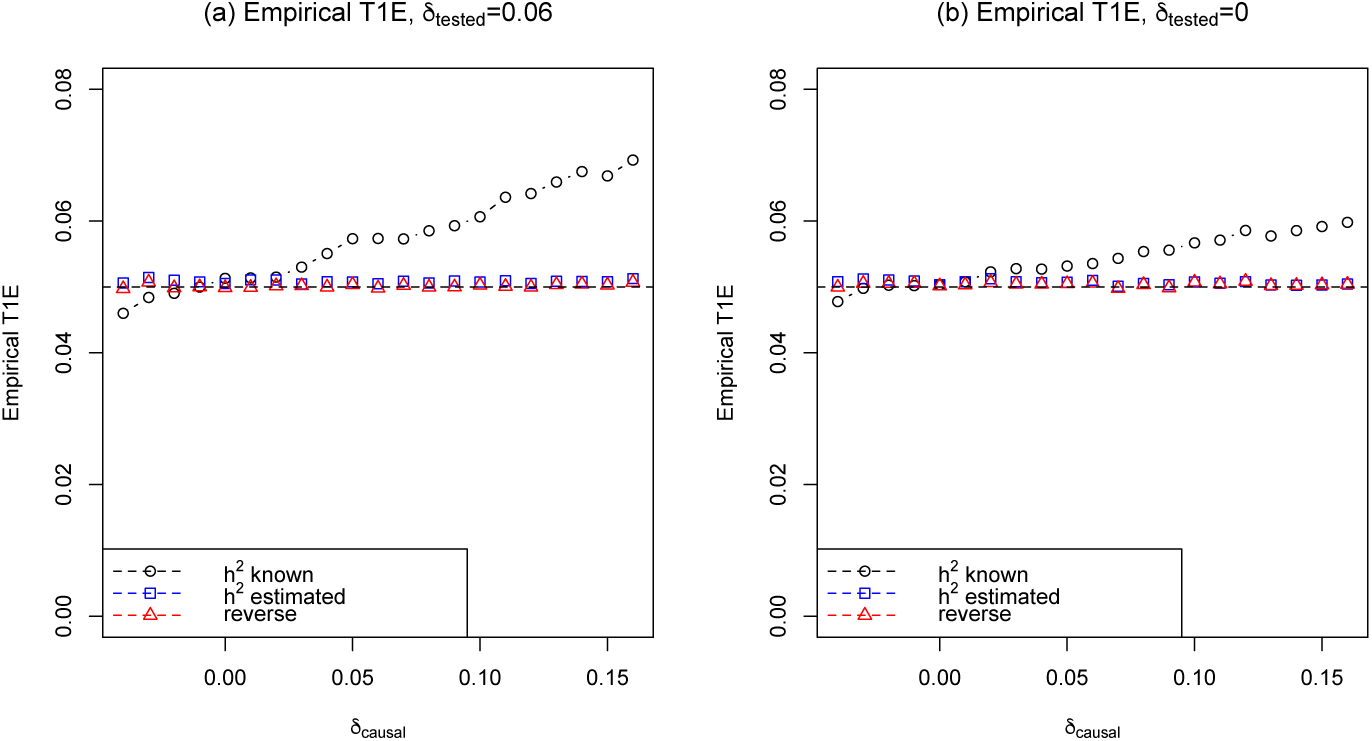
Empirical type I error rate of *T*^LMM^ based on the linear mixed-effect model (4) and *T*_reverse_ based on the proposed ‘reverse’ regression model (6), against *δ*_causal_. (a) When *G*_tested_ of tested SNPs are in HWD with *δ*_tested_ = 0.06. (b) When *G*_tested_ of tested SNPs are in HWE with *δ*_tested_ = 0. The true heritability of the phenotype is *h*^2^ = 0.5, the minor allele frequencies *p*_causal_ = *p*_tested_ = 0.2, and 10, 000 independent replicates of phenotypes and genotypes for 65 sibling pairs were simulated for each *δ*_causal_ value. Black circles are for *T*_LMM_ using the true *h*^2^ = 0.5, blue circles are for *T*_LMM_ while estimating *h*^2^ (results of *ĥ*^2^ shown in Figure 3), and red triangles are for *T*_reverse_.

In most practical implementations of the linear mixed-effect model (4), *h*^2^ is treated as a nuisance parameter, and no type I error issue has been reported. Indeed, when *h*^2^ was estimated in our simulation study, the size of *T*_LMM_ was correct (blue squares in Figure 2). However, in this situation, the impact of HWD is now on the estimation of *h*^2^. Specifically, we applied the LMM (4) to the same simulated data as above but assumed *h*^2^ to be unknown. Figure 3 clearly shows that *ĥ*^2^ is downward biased when *δ*_causal_ *<* 0, and upward biased if *δ*_causal_ *>* 0. The bias can be substantial. For example, when *δ*_causal_ = 0.10, the estimated heritability *ĥ*^2^ is centred at 0.78 as compared to the true value of 0.5.

**Figure 3:**
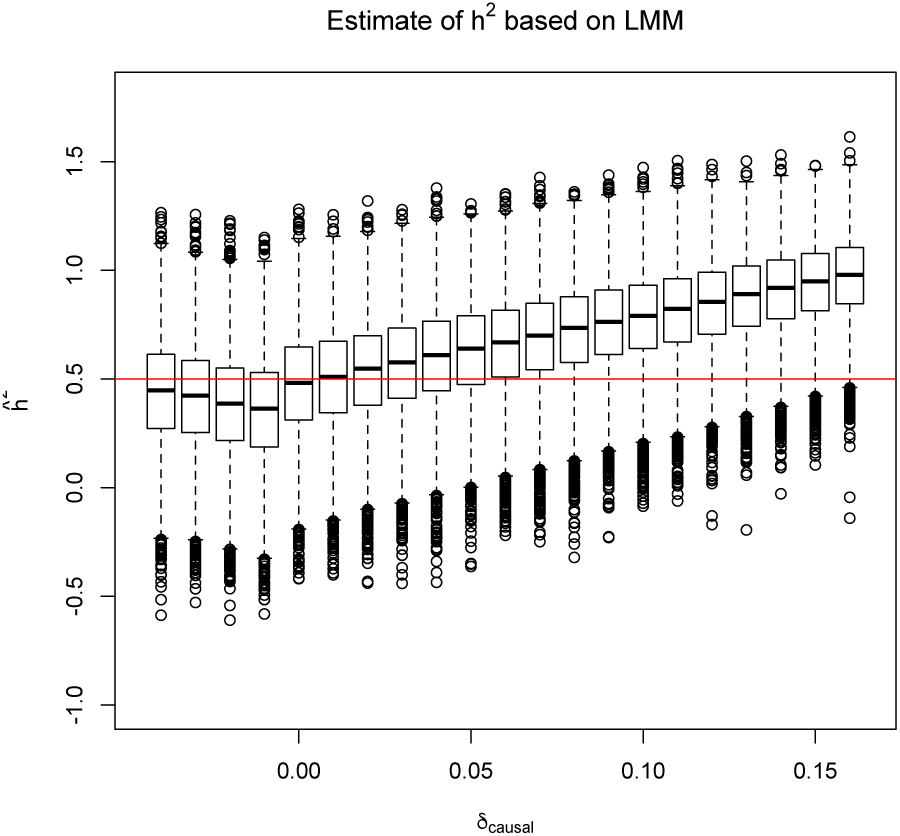
Box-plots of *ĥ*^2^, estimated from the linear mixed-effect model (4) against *δ*_causal_. The true heritability of the phenotype is *h*^2^ = 0.5. The minor allele frequencies *p*_causal_ = *p*_tested_ = 0.2, and 10, 000 independent replicates of phenotypes and genotypes for 65 sibling pairs were simulated for each *δ*_causal_ value. The empirical type 1 error rates are shown in Figure 2 as blue circles.

In Figure 3, it is notable that when *δ*_causal_ *>* 0.1, *ĥ*^2^ *>* 1. Since *h*^2^ is the proportion of variance in *Y* explained by additive genetic variation, 0 *≤ h*^2^ *≤* 1 by definition. However, if *δ*_causal_ ≠ 0, *ĥ*^2^ based on the LMM, without additional truncation, is a biased estimate of *h*^2^ by a factor of 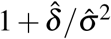.

### 3.3 Estimation of minor allele frequency

Here we compare the best linear unbiased estimator (BLUE) by McPeek et al. (2004) with the proposed ‘reverse’ regression estimator, in the presence of sample correlation and Hardy–Weinberg disequilibrium. We chose the parent-child relationship for this study. Note that even though the kinship coefficient of a parent-child pair is identical to that of a sib-pair, Σ_*δ*_ is not the same for these

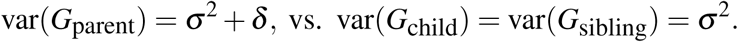

For a parent-child pair,

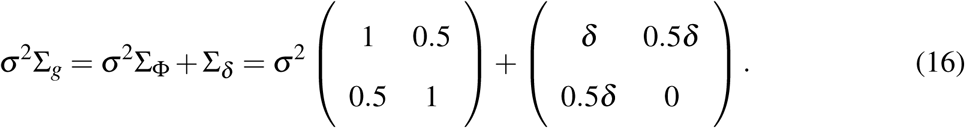

Let 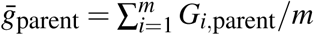 and 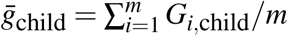, where *m* is the number of independent families, the minor allele frequency estimates are

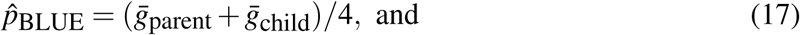

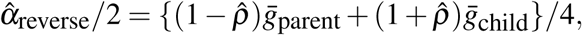

Where

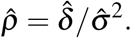

It is easy to show that 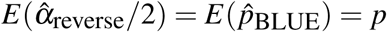 regardless of the value of *ρ*.

In equation (17), 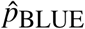 always weighs 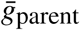 and 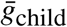 equally. For the proposed estimator 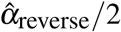 if 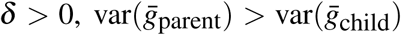. Accordingly, because *ρ* > 0, 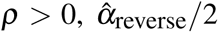 would weigh 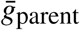 less than 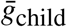 Conversely, if *δ* < 0, 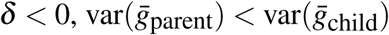, and because *ρ* < 0, 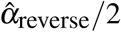 would weigh 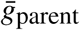 more than 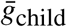 Therefore, variance of 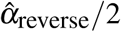 is smaller than that of 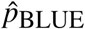

To confirm this analytical insights numerically, we simulated 1, 000 parent-child pairs, where the parents’ genotypes were out of HWE, with *δ*_parent_ = 0.06 and *p*_parent_ = 0.2. A total of 50, 000 independent replicates were simulated, and the corresponding minor allele frequency estimates by the two methods are shown in Figure 4(a). To demonstrate the empirical standard error of the estimates, we first randomly split the 50, 000 estimates into 100 even sets, then calculated the sample standard error based on the 500 estimates from each set. The corresponding 100 standard error estimates are shown in Figure 4(b). Results clearly demonstrate the estimation efficiency of the proposed 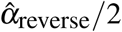 as compared with 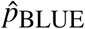 in the presence of HWD for parent-child pairs.

**Figure 4:**
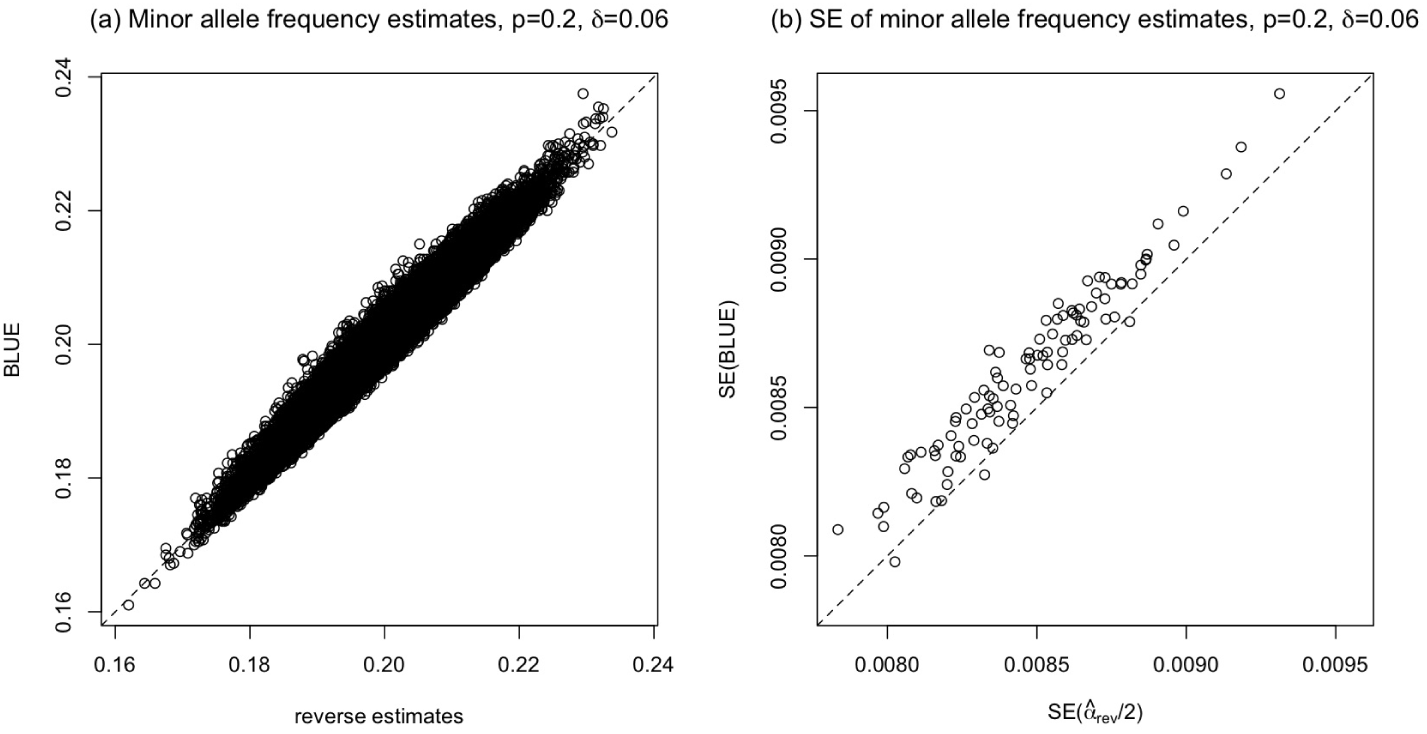
Results of the minor allele frequency estimation by BLUE and the proposed estimator (17) for simulated parent-child data. The proposed estimator is a special case of 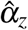 of (15), derived from the proposed ‘reverse’ regression without covariate *Z*. (a) Minor allele frequency estimates based on 50, 000 independent replicates. (b) Empirical standard error of the minor allele frequency estimates, based on 100 independent sets of 500 estimates. Each minor allele frequency estimation was based on 1, 000 parent-child pairs, *p*_parent_ = 0.2 and *δ*_parent_ = 0.06 (offspring data are determined by parental data), with 500 × 100 independently simulated replicates.

## 4 Discussion

In this note, after reviewing a catalogue of genetic association tests we proposed a ‘reverse’ regression model, treating genotype *G* as the response variable and phenotype *Y* as a covariate. The proposed method is robust and flexible, and it can (i) analyze both binary and continuous outcomes, (ii) adjust for additional covariate effects, (iii) handle dependent samples, and (iv) incorporate a factor *δ* to adjust for potential departure from Hardy–Weinberg equilibrium. Furthermore, the proposed *T*_reverse_ unifies many existing genotype-based tests, including *T*_LMM_ based on the linear mixed-effect model, quasi-likelihood score test for binary traits *T*_QL-binary_ (Thornton and McPeek, 2007), generalized quasi-likelihood score test *T*_GQLS_ (Feng et al., 2011; Feng, 2014), and *T*_MASTOR_ (Jakobsdottir and McPeek, 2013). Finally, the proposed regression framework can also estimate allele frequency while adjusting for covariates and departure from HWE. Table 1 provides a summary of the method comparison.

**Table 1:**
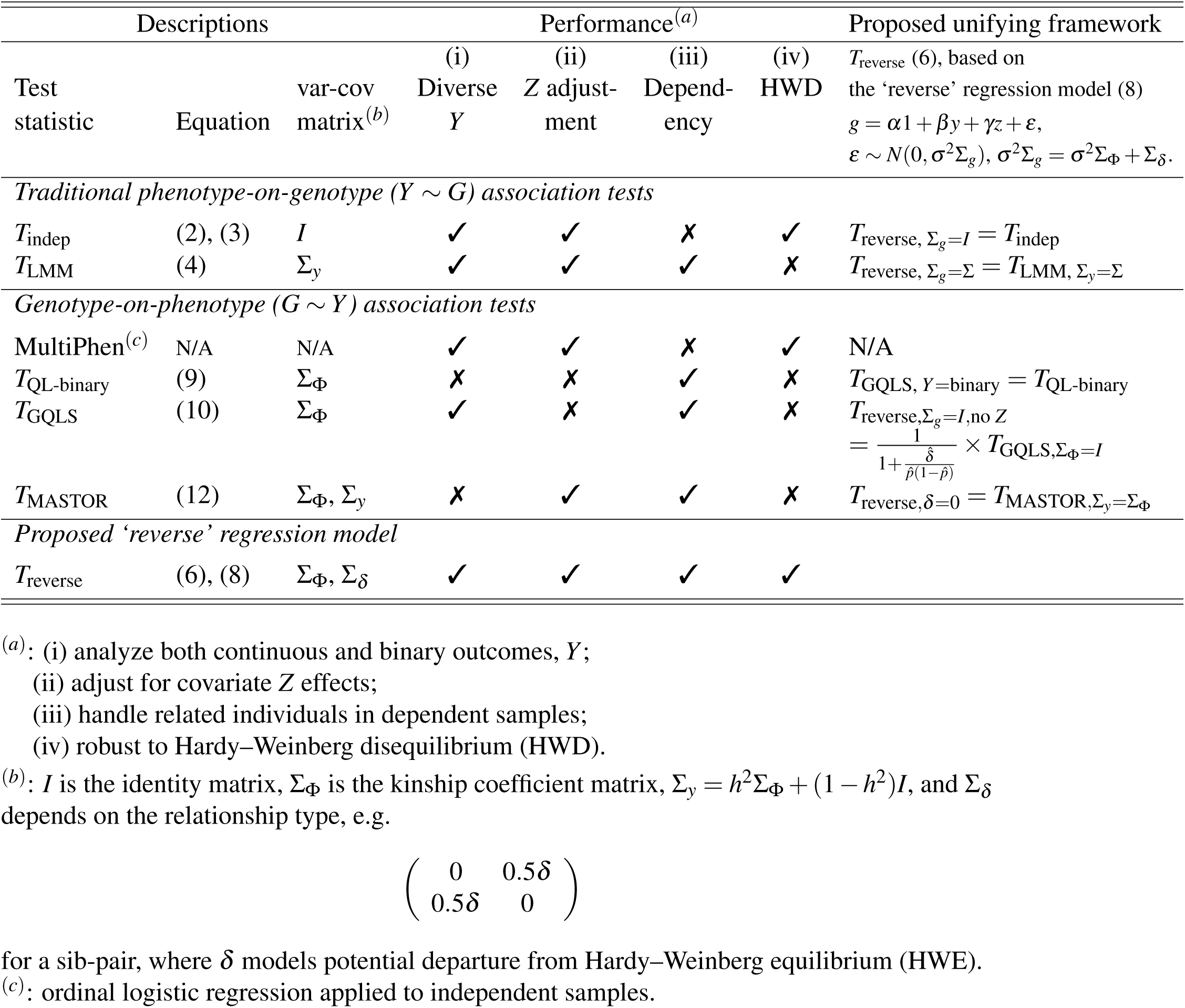
Comparison of genotype-based association tests.

In the ‘reverse’ regression model, we treated the discrete genotype data *G* as continuous, because the method is purposed for association testing and earlier work has shown that, under some regularity conditions, the score test statistic for the exponential family is identical (Chen, 1983). Indeed, for the simple cases of independent samples, no covariates, or HWE, desirably the proposed test numerically coincides with a number of existing methods as shown in Section 2.

Departure from HWE generally falls into two categories: disequilibrium at the causal SNP(s) and disequilibrium at the tested SNP. If HWE does not hold at the causal SNP(s), models that rely on σ_*y*_ (e.g. the linear mixed-effect model and MASTOR) may have inflated type I error rate if *δ*_causal_ > 0, or deflated type I error rate if *δ*_causal_ < 0, assuming the true heritability *h*^2^ is known; if *h*^2^ is estimated based on the data, the estimate is then biased if *δ* ≠ 0. If Hardy–Weinberg disequilibrium occurs at the tested SNP, models that assumes 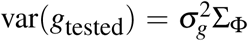 (e.g. quasilikelihood score test and MASTOR) may be sensitive to *δ*_tested_ ≠ 0.

At a tested SNP, because the proposed ‘reverse’ framework is conditional on *Y*, the variance-covariance matrix only concerns *G*_tested_, i.e. ∑ _*g*_. The modelling and estimation of ∑ _*g*_ can account for potential departure from HWE through ∑ _*δ*_, in addition to genetic correlation as captured by the kinship coefficient matrix of ∑ _Φ_, resulting in a more robust association test for related individuals.

We demonstrated the type I error issue of the linear mixed-effect model, in the presence of HWD and assuming the true heritability is known, using data consists of related individuals only. In practice, this issue diminishes if the sample includes a large number of independent individuals or the magnitude of HWD is small. Nevertheless, the analytical framework presented here can be an useful alternative that not only account for potential HWD but also unifies a number of existing tests. Even if HWD is of no concern, the proposed framework generalizes the MultiPhen approach (O’Reilly et al., 2012) to jointly analyze multiple phenotypes using related individuals. Further, we demonstrated that when *h*2 is treated as unknown, its estimation can be biased and often upwardly (if 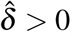) as seen in the CF application. This insight offers a possible alternative interpretation of the ‘missing heritability’ issue (Maher, 2008).

In practice, SNPs out of HWE are typically not analyzed due to concerns for low genotyping quality. However, the issue identified here remains relevant: HWE quality control screening uses a *p*-value threshold in the range of 10^−8^ (Consortium et al., 2007), and this practice itself can be called into question because a truly associated SNP is likely to be out of HWE (Turner et al., 2011; Ryckman and Williams, 2008). The potential of using the proposed framework to jointly test *δ* and *β* to increase the power of association testing is of future research interest.

For genetic epidemiological interpretation, it may appear that the traditional *Y* -on-*G* regression approach is more intuitive. But, as pointed out by Schaid et al. (2013) in a different setting, the retrospective alternative that treats *G* as random while conditioning on *Y* “overcomes the problem of modelling the ascertainment process, which would be particularly challenging for highly enriched pedigrees.” How to model gene-environment interaction using the proposal ‘reverse’ regression framework, however, remains an open question.

## Acknowledgements

We thank Dr. Lisa J. Strug and her lab for providing the cystic fibrosis genotype data. This research is funded by the Natural Sciences and Engineering Research Council of Canada (NSERC, RGPIN-04934 and RGPAS-522594), and the Canadian Institutes of Health Research (CIHR, MOP-310732) to LS. LZ is a trainee of the CIHR STAGE (Strategic Training in Advanced Genetic Epidemiology) training program at the University of Toronto.

